# *In silico* and *in vitro* Demonstration of Homoharrintonine’s Antagonism of RBD-ACE2 Binding and its Anti-inflammatory and anti-thrombogenic Properties in a 3D human vascular lung model

**DOI:** 10.1101/2021.05.02.442384

**Authors:** Shalini Saxena, Kranti Meher, Madhuri Rotella, Subhramanyam Vangala, Satish Chandran, Nikhil Malhotra, Ratnakar Palakodeti, Sreedhara R Voleti, Uday Saxena

## Abstract

Since 2019 the world has seen severe onslaught of SARS-CoV-2 viral pandemic. There is an urgent need for drugs that can be used to either prevent or treat the potentially fatal disease COVD-19. To this end, we screened FDA approved antiviral drugs which could be repurposed for COVID-19 through molecular docking approach in the various active sites of receptor binding domain (RBD). The RBD domain of SARS-CoV-2 spike protein is a promising drug target due to its pivotal role in viral-host attachment. Specifically, we focussed on identifying antiviral drugs which could a) block the entry of virus into host cells, b) demonstrate anti-inflammatory and/or anti-thrombogenic properties. Drugs which poses both properties could be useful for prevention and treatment of the disease. While we prioritized a few antiviral drugs based on molecular docking, corroboration with *in vitro* studies including a new 3D human vascular lung model strongly supported the potential of **Homoharringtonine**, a drug approved for chronic myeloid leukaemia to be repurposed for COVID-19. This natural product drug not only antagonized the biding of SARS-CoV-2 spike protein RBD binding to human angiotensin receptor 2 (ACE-2) protein but also demonstrated for the first time anti-thrombogenic and anti-leukocyte adhesive properties in a human cell model system. Overall, this work provides an important lead for development of rapid treatment of COVID-19 and also establishes a screening paradigm using molecular modelling and 3D human vascular lung model of disease to identify drugs with multiple desirable properties for prevention and treatment of COVID-19.

## 1. Introduction

SARS-CoV-2 virus has devastated the world since late 2019. The virus when left untreated leads to COVID-19 disease and is often fatal. The disease begins with entry of the virus into the human host cells. Coronaviruses (CoVs), including SARS-CoV, MERS-CoV, and SARS-CoV-2, are cytoplasmically replicating, positive-sense, single-stranded RNA viruses with four structural proteins *i.e.* Spike protein (S), envelope (E) protein, membrane (M) protein, and nucleocapsid (N) protein [1].

This entry of SARCS-CoV-2 into host cells is mediated by a specific receptor binding domain (RBD) on the surface of spike protein. The entry of the virus into the host cell requires the spike or S-protein to be cleaved in two steps. The first step is the binding of the virus to the ACE-2 protein and this is accomplished by the cellular proteases acting at the region between S1 and S2, namely, the transmembrane serine protease TMPRSS2. S1 and S2 subunits are responsible for viral-receptor binding and virus-host cell membrane fusion, respectively. The S1 subunit bears the receptor-binding domain (RBD) on its C-terminal domain. The RBD itself contains the receptor-binding motif (RBM), which actually comes into contact with the carboxypeptidase domain of the ACE-2 molecule. The RBD engages with a counter receptor on human cells called angiotensin receptor 2 (ACE-2) and gains entry. Once inside the human cells, it uses host machinery to replicate itself. Thus, antagonism of binding of RBD to ACE-2 could be a preventive strategy for this infection. Generally, the S protein plays a crucial role in eliciting the immune response during disease progression also. Thus S protein RBD domain, if inhibited, should prevent viral attachment, fusion and entry, and thus make it an attractive target over other targets [2].

Several human cells express ACE-2 on their surface, lung cells, intestinal cells, endothelial cells, kidney cells among a few that have been clearly demonstrated in humans. These receptors/cells serve as chief mediators of viral entry and dysfunction of that particular organ. The impact of viral attachment and entry has been best studied in lungs. Five clear steps have been delineated based on human autopsy and lung biopsy studies

a. Viral entry into lung epithelial cells.
b. Activation of surrounding vascular capillary endothelium which is abundant in lungs.
c. The activation of endothelial cells by virus leading to recruitment of inflammatory cells such as monocytes into the lung.
d. Uncontrolled production of inflammatory cytokines “cytokine storm” further exaggerating lung dysfunction.
e. A prothrombogenic environment which promotes blood clotting in lung capillaries.

All these events translate into a hostile and highly inflammatory environment in the lungs. There is a rapid decline in lung function due to lung cell death, fibrosis and hypoxia. Most other organs also fail in due time due to direct viral intervention and or hypoxia due to lack of oxygen supply by blood.

But such events also provide choke points for therapeutic intervention. For example, a drug that combines inhibiting the binding of virus to human cells and also possess anti-inflammatory and anti-thrombogenic properties may be more desirable than a drug with just anti-inflammatory property or entry antagonism alone. **Table-1** lists our proposed wish list of properties that may be desirable for a drug to be repurposed for prevention and treatment.

**Table-1:** List of properties that are desirable for repurposing of a drug for COVID-19

| Property | Desired Properties |
| --- | --- |
| History of extensive human safety | Yes |
| Ease of availability and cost globally | Yes |
| Block viral entry into human cells | Yes |
| Block inflammation (cytokine secretion and monocyte adhesion) | Yes |
| Reduce excess blood clotting | Yes |

With this in mind we started a search for drugs with multiple therapeutic properties. While there are now validated animal models of this disease, whether they capture the complete pathogenesis seen in humans remains to be seen. In this respect, it has been proposed by others that current animal models mainly capture the early and mild form of the disease [3]. Also, animal models require Biosafety level BSL 3 or 4 lab facilities and the models have low drug screening throughout. In order to replicate above stated features of human COVID-19, we created a 3D bioprinted human vascular cell model. In this model, immortalized human lung epithelial cells (A549) are bioprinted in layers along with human endothelial cells interspersed with cell growth matrix bioink. The model can be used to study lung cell function, cytokine secretion, endothelial dysfunction by measuring secretion of thrombogenic factors. Finally, we can also mimic monocyte adhesion by adding monocytes to the 3D bioprinted system. So, using one model we are able to screen for several features seen in human disease.

We chose a few selected antiviral drugs with demonstrated efficacy against other viruses such MERS and SARS. The idea behind this was that since they have antiviral activity to begin with, we are likely to guarantee ourselves of one potential therapeutic activity. Secondly, since they are all drugs approved for human use, they can be rapidly deployed for COVD-19 treatment. Since the entry of virus initiates the pathological cascade of evets, we first rank ordered the drugs based on their ability to block the binding of RBD to ACE-2 using molecular docking studies. After that we systematically tested them for a) inhibition of binding of RBD to ACE-2 in a cell free ELISA assay to demonstrate a direct effect (without the background of cell based assays confounded my drug metabolism penetration and drug solubility in cell media) b) Effect of drugs on secretion of IL-1β by the lung cells c) Effect of drugs on secretion of thrombomodulin, a thrombin activity modulating factor involved in clotting and finally d) impact of drugs on adhesion of monocytes to the vascular cells in 3D model. We find that Homoharringtonine, a natural product inspired drug approved for human use, possess many of the properties desirable for repurposing a drug for COVID-19.

## 2. Material and Methods

### 2.1 Computational studies

Cresset Flare software was used for molecular docking studies against the spike protein SARS-CoV-2 (http://www.cresset-group.com/flare/) [4].

#### 2.1.1 Ligand preparation

The 2D structures of all the antiviral drugs were downloaded from Drugbank database [5] and prepared using Flare software. Atom force field parameterization was assigned and hydrogen atoms were added in the structure. Further, energy minimized was done for all the drugs, nonpolar hydrogen atoms were merged, and rotatable bonds were defined. Later, ligand minimization has been carried out in Flare by Minimize tool by using Normal calculation methods.

#### 2.1.2 RBD structure preparation

The RBD of spike glycoprotein SARS-CoV-2 is used for present study. It is clear that spike proteins represent potential targets for anti-SARS-CoV-2 drugs hampering the interaction between human ACE-2 and the viral RBD will block the entry of the virus into the human cells. The 3D structure of RBD binds to ACE-2 receptor (PDB ID: 6M0J) were downloaded from Protein Data Bank (PDB) (https://www.rcsb.org) [6]. The protein has two chain A and E, the A chain has ACE-2 receptor and E chain has RBD domain. The RBD domain has been save in to PBD format for further studies. The target protein preparation was carried out in Flare software with default settings. Missing residues, hydrogen's and 3D protonation were carried out on the target protein. Protein minimization has been carried out in Flare by Minimize tool by using Normal calculation methods.

#### 2.1.3 Computational analysis of binding sites for RBD

Binding site was generated Accelrys Discovery Studio visualizer 3.5 (Copyright© 2005-12, Accelrys Software Inc.) to explore potential binding sites of the RBD protein using receptor cavities tools. Based on a grid search and “eraser” algorithm, the program defines where a binding site is. The binding sites were displayed as a set of points (point count) and the volume of each of cavity was calculated as the product of the number of site points and the cube of the grid spacing. Volume of each site were calculated and further saved and exported in to Flare for advance analysis.

#### 2.1.4 Molecular docking

All the 23 antivirals were docked in the active site of RBD by using Flare docking module of Cresset software. All the compounds were subjected to docking using Lead Finder (LF) and the predicted binding poses were analysed [7] docking analysis and visualization of hits led to the identification of 7 drugs based on rank score (RS), and binding energy (∆G) that bind selectively to the RBD site-1 and site-2.

### 2.2 Experimental studies

#### 2.2.1 RBD-ACE-2 biochemical assay

The above assay was performed using SARS-CoV-2 sVNT ready to use kit by Genscript, which is a competition ELISA, mirroring the viral neutralization process. In the first step, a mixture of horse radish peroxidase-RBD (HRP-RBD) and controls/drugs (drugs at the concentrations of 50μM, 100μM and 200μM) were incubated at 37° C for an hour to allow the binding of drugs to HRP-RBD. Following the incubation, these mixtures were added to a capture plate, which was a 96 well microplate coated with human ACE-2 receptor to permit the binding of any unbound HRP-RBD and the ones bound to drugs to ACE-2 receptor. After incubating the microplate at 37° C for an hour, plate was washed four times using wash buffer in order to remove any unbound circulating HRP-RBD_ drug complexes. Washing step was followed by addition of a colour substrate; tetramethyleneblue (TMB), turning the colour to blue. The reaction was allowed to run for 15 minutes followed by the quenching using stop solution turning the colour from blue to yellow. This final solution was read at 450 nm. The absorbance of the sample is inversely proportional to the inhibition of RBD’s binding to the human ACE-2 by the drug.

#### 2.2.2 IL-1ẞ secretion assay

Human IL-1ẞ ELISA was performed to detect the presence of IL-1ẞ which is a key mediator of inflammatory response. The ELISA was performed using Human IL-1ẞ high sensitivity kit sold by Invitrogen. The sample for our ELISA consisted of cell culture media (referred to as sample hereafter) collected after treating the cells with drugs at 100uM. Samples were added to the microplate precoated with human IL-1ẞ antibody which captures the IL-1ẞ present in the samples. A secondary anti-human IL-1ẞ antibody conjugated to biotin was added to the plate. Following an overnight incubation, microplate was washed six times using wash buffer in order to remove any unbound biotin conjugated anti-human IL-1ẞ antibody. Streptavidin-HRP was then added which binds to biotin conjugated antibody and the plate was incubated at room temperature on a shaker for an hour. After the incubation, plate was washed again following the same process as the previous wash step and an amplification reagent I was added to the wells. Following the incubation of 15 minutes and a wash, amplification reagent II was added. After incubation of half an hour in dark and a wash step later, a substrate solution was added turning the colour to blue. The reaction was terminated using a stop solution (turning the colour from blue to yellow) after 15-20 minutes. This final solution was read at 450nm. The OD of the sample is directly proportional to the amount of human IL-1ẞ present in it.

#### 2.2.3 Human Thrombodulin/BDCA-3 Immunoassay

Human thrombodulin ELISA was performed using a ready to use kit by R and D systems. Samples (media collected from our 3D vascular lung system) were added to a microplate pre-coated with monoclonal antibody specific for human thrombodulin. This was followed by incubation period of two hours (allowing any thrombodulin present in the sample to bind to the monoclonal antibody) the plate was washed with wash buffer four times. After washing away any unbound thrombodulin, an enzyme-linked monoclonal antibody specific for human thrombodulin was added to the plate. Another wash step was performed in order to remove any unbound antibody-enzyme reagent after completion of two hours of incubation. A colour substrate was then added turning the colour to blue. Reaction was quenched after about 15-20 minutes using a stop solution which turned the colour from blue to yellow. Absorbance was read at 450 with wavelength correction set to 540 nm or 570 nm.

#### 2.2.4 Cell culture

For the 3D vascular lung model three types of cell were grown:

1. A549 cells were grown at 37° C in the growth medium DMEM (HIMEDIA #Cat No- AT007) supplemented with 10% (v/v) fetal bovine serum under the atmosphere containing 5% CO2. Cells were subcultured after reaching 80-90% confluence.
2. HUVEC cells were grown at 37° C in the Endothelial cell basal medium-2 (LONZA #Cat No-CC-3156 and CC-4176) under the atmosphere containing 5% CO2. Cells were subcultured after reaching 80-90% confluence.
3. HL60 cells were grown at 37° C in the growth medium RPMI (HIMEDIA #Cat No- AT028) supplemented with 10% (v/v) fetal bovine serum under the atmosphere containing 5% CO2. Cells were subcultured after reaching 80-90% confluence.

#### 2.2.5 3D-bioprinting

In the 3D-vascular lung model four layers were bioprinted, first layer was collagen layer, (30μl of Rat tail collagen) was added to each well in 96 well plate and incubated for 1 hour in CO2 incubator at 37°C. After incubation, once the collagen is solidified A549 cells flask which has reached 80-90% confluence were trypsinized and cells were counted with the help of haemocytometer. Cell suspension was loaded into bioprinter syringe and 5×10^3^ A549 cells were printed in each well of 96 well plate and the cells were incubated for 48 hours. After 48 hours of incubation, previous A549 media was removed carefully and again 30μl of Rat tail collagen coating solution was added to each well in 96 well plate and incubated for 1 hour in CO2 incubator at 37°C. After incubation, once the collagen is solidified HUVEC cells flask which has reached 80-90% confluence was trypsinized. Cell suspension was loaded into syringe and 5×10^3^ HUVEC cells were printed in each well of 96 well plate with help of 3D bioprinter and the cells were incubated for 48 hours.

After 48 hours of incubation the media was removed and the final selected drugs from virtual screening were added with LPS at 0.5 μg/ml and without LPS at the desired concentration. Later, cells were incubated overnight. Next day the drugs were removed and the cells were washed once with media. Endothelial cells were fixed with 4% paraformaldehyde for 3 minutes at 4°C. After fixing with paraformaldehyde, the cells were again washed with media.

MTT stained HL60 monocytic cells 1× 10^4^ cells were added to each well in 96 well plate and incubated the plate for one hour and the wells were then washed twice with the media and pictures were taken of all the wells and the bound HL 60 cells were counted with the help of ImageJ, an open access software platform.

## 3. Results and Discussion

### 3.1 Identification of active sites (RBD)

Currently, there are no specific effective antiviral treatments for COVID-19, although most of the COVID-19 patients have mild or moderate courses, up to 5%-10% can have severe, potentially life threatening course. Thus, there is an urgent need for effective drugs [8]. Our *in silico* strategy helps us to find new repurposed antiviral drugs that can attach to residues at the site of binding of the RBD to the ACE-2. In the current study we had discovered several potential binding sites for molecules that can occupy such druggable pockets so as to inhibit virus-ACE-2 binding *in vitro*. The X-ray model of the RBD was used to identify 3 possibly druggable pockets where drugs might bind. The active site volume and binding surface area of three pocket is representation in **Table-2**. Site-1 has the volume of 143.7 & site-2 has the active site volume of 109.4, while the last site-3 has active site volume of 87.46. Site-3 was too small to accommodate ligands, so it could not be potential pocket to accommodate antagonists. Based on size of active site cavity site-1 & site-2 has been selected for further docking studies.

**Table-2:** Active sites of RBD domain of spike protein.

| Site | Binding surface area ($\text{\AA}^2$) | Active site volume ($\text{\AA}^3$) |
| --- | --- | --- |
| S1-1 | 673.6 | 143.7 |
| S1-2 | 668.3 | 109.4 |
| S1-3 | - | 87.46 |

### 3.2 Molecular Docking and Interaction Analysis in RBD Domain

Docking analysis and visualization of selected antivirals led to the identification of 7 drugs that bind selectively to the both the RBD sites. Total 7 drugs were identified based on rank score (RS), and binding energy (∆G). These include Homoharrigtonine, Triparanol, Lopinavir, Ritonavir, Astemizole, Amodiaquine, and Fluspirilene. Our goal was to find those molecules which have great binding affinity towards both the active sites of RBD domain.

Detail binding analysis of selected drugs towards active site of spike protein SARS-CoV-2 was studied in detail. Interaction analysis of drugs with spike protein SARS-CoV-2 (RBD) were carried out to identify the compound having highest binding affinity with target protein. Active site-1 composed of Arg454, Phe456, Arg457, Lys458, Glu465, Arg466, Asp467, Ile468, Ser469, Glu471, Thr473, Gln474 and Pro491 amino acid residues, while the site-2 composed of Leu335, Cys336, Pro337, Phe338, Gly339, Trp436, Phe342, Asn343, Val362, Ala363, Asp364, Val367, Leu368, Ser371, Ser373 and Phe374 amino acid residues, as shown in **Figure-1**. Active site-1 is more hydrophobic in nature as compare to active site-2.

**Figure-1:**
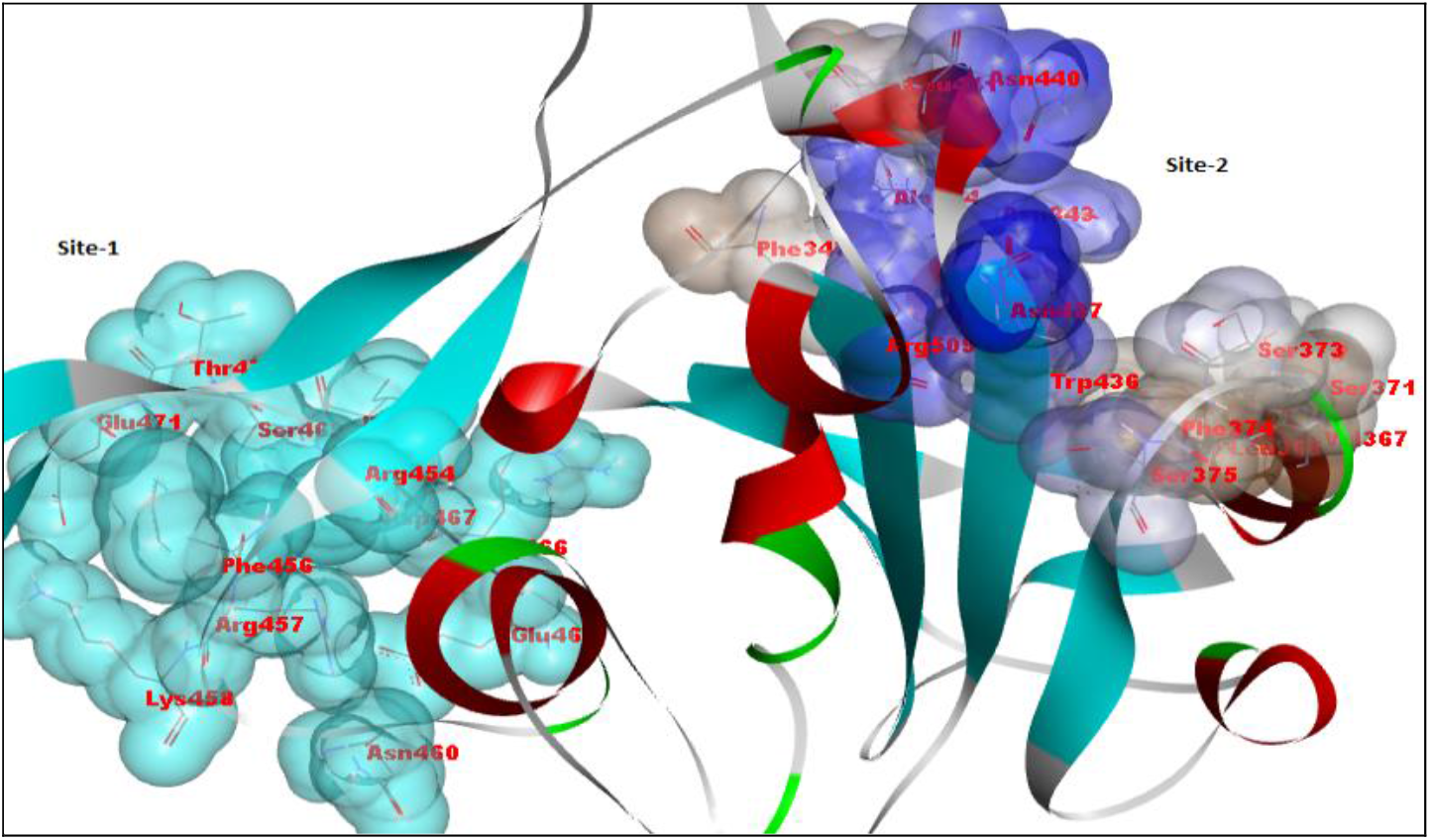
Active sites of RBD domain of S1 protein.

The LF rank score is an indicator of the binding affinity of protein-ligand complex. The LF rank for selected drugs in the pockets site-1 & site-2 is described in **Table-3** respectively. The binding orientation for each selected drugs having the least LF rank score, more negative LF rank score represent the better affinity of the drugs against target SARS-CoV-2 S-protein. Among the docking studies performed on drugs, 7 drugs had effective binding interactions with RBD domain of SARS-CoV-2 S-protein in both the active sites. The LF rank score values were in the range of −4.1 to −10.5 in the active site 1, while in active site 2 the LF rank score range from −4.0 to −10.6 for all the selected 23 antiviral drugs. The ∆G provide an accurate estimate of protein-ligand binding energy, on the assumption that the pose is correct. For the selected drugs the ∆G is range from −3.2 to −9.7 in site-1, while in site-2 the ∆G is range from −4.8 to −9.7.

**Table-3:** Docking score of 23 antiviral drugs for SARS-CoV-2 Spike RBD in site-1 and −2.

|  | Spike Protein S1-1 |  | Spike Protein S1-2 |  |
| --- | --- | --- | --- | --- |
| Drug Name | Rank score | $\Delta G$ | Rank score | $\Delta G$ |
| **Lopinavir\*** | **-10.5** | **-6.3** | **-9.2** | **-8.1** |
| **Ritonavir\*** | **-10.3** | **-9.7** | **-9.4** | **-9.7** |
| **Amodiaquine\*** | **-10.2** | **-6.7** | **-9.4** | **-5.0** |
| Anisomycin | -7.9 | -5.3 | -8.2 | -7.4 |
| **Astemizole\*** | **-10.1** | **-6.0** | **-10.6** | **-7.6** |
| Benzotropine | -4.7 | -5.6 | -4.2 | -6.0 |
| Chloroquine** | -6.1 | -3.2 | -3.4 | -7.8 |
| Chlorphenoxamine | -4.3 | -5.4 | -4 | -5.8 |
| Cycloheximide | -8.7 | -7.6 | -6.7 | -7.3 |
| Emetine | -4.7 | -8.1 | -4 | -6.9 |
| Fluphenazine | -8.8 | -8.6 | -8.1 | -6.5 |
| **Fluspirilene\*** | **-10.5** | **-5.7** | **-8.5** | **-7.3** |
| Gemcitabine | -6.5 | -4.1 | -6.2 | -4.8 |
| **Homoharringtonine\*** | **-9.0** | **-7.4** | **-8.2** | **-7.5** |
| Hydroxychloroquine | -8.5 | -7.2 | -8.5 | -8.1 |
| Promethazine | -8.4 | -4.9 | -7.7 | -5.8 |
| Tamoxifen | -4.1 | -5.4 | 4 | -6.2 |
| Terconazole | -5.3 | -8.0 | -4.6 | -8.4 |
| Thiethylperazine | -8.4 | -7.2 | -8.2 | -7.2 |
| Thiothixene | -8.4 | -8.1 | -8.3 | -8.7 |
| Toremifene** | -4.1 | -5.9 | -3.7 | -6.9 |
| Triflupromazine | -7.9 | -5.7 | -7 | -6.3 |
| **Triparanol \*** | **-9.4** | **-7.5** | **-9.6** | **-8.4** |

\*Highlighted drugs were selected for biological evaluations.
\*\* Lowest scoring drugs used for biological evaluations

Out of 23 antivirals, we selected 7 drugs for *in vitro* biological evaluation based on rank score and ∆G along with their interactions with the protein. We also selected two low scoring drugs (Toremifene and Chloroquine) for *in vitro* biological evaluation to confirm ours *in silico* studies.

As shown in **Figure-2**, the graph represents the correlation between rank score and binding energies of all the selected drugs in RBD site-1 and site-2. All the drugs have more or less similar range of score and binding energies. Also, the selected drugs well occupied the active site cavity (S1-1 and S1-2).

**Figure-2:**
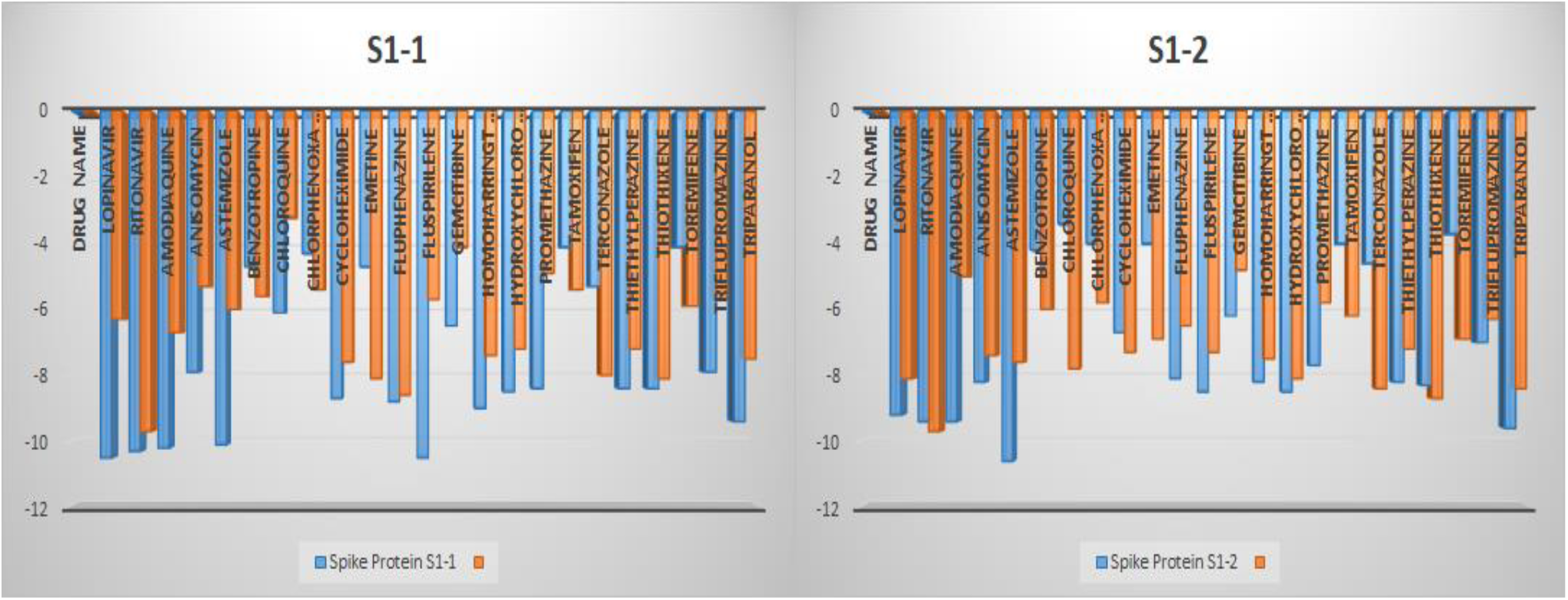
Rank score and binding energies of antiviral drugs in RBD domains.

Detail interaction analysis was performed for the selected drugs which have good binding affinity towards RBD domain. The identified Lopinavir and Ritonavir drugs are fixed-dose combination antiretroviral medication for the treatment and prevention of HIV/AIDS. In our study, both the drugs had good binding with the RBD protein. The docking orientation of Lopinavir and Ritonavir in site-1 & site-2 is represented in **Figure-3** and **-4**, respectively. Lopinavir has good binding affinity towards RBD having rank scores of −10.4 in site-1, and - 10.6 in site-2, while Ritonavir has rank score of −10.3 and −9.8 in both the sites. In site-1, the drug is stabilized by five hydrogen bonding interactions with Arg457, Lys458, Asn460 and Asp467 amino acid residues. Apart from this the drug is making various hydrophobic interactions with Glu471, Phe456, Ile468, Ser469 and Glu465. It has strong *cation-pi* interaction with Asp467 and Arg454. In the active site 2 the drug is making hydrogen bond with Asn343 and Trp436 amino acid residue. The drug is making a close contact with various hydrophobic interactions with Phe374, Ser371, Phe342, Ala344, Ile441, Asn440, Phe338 and Gly339.

**Figure-3:**
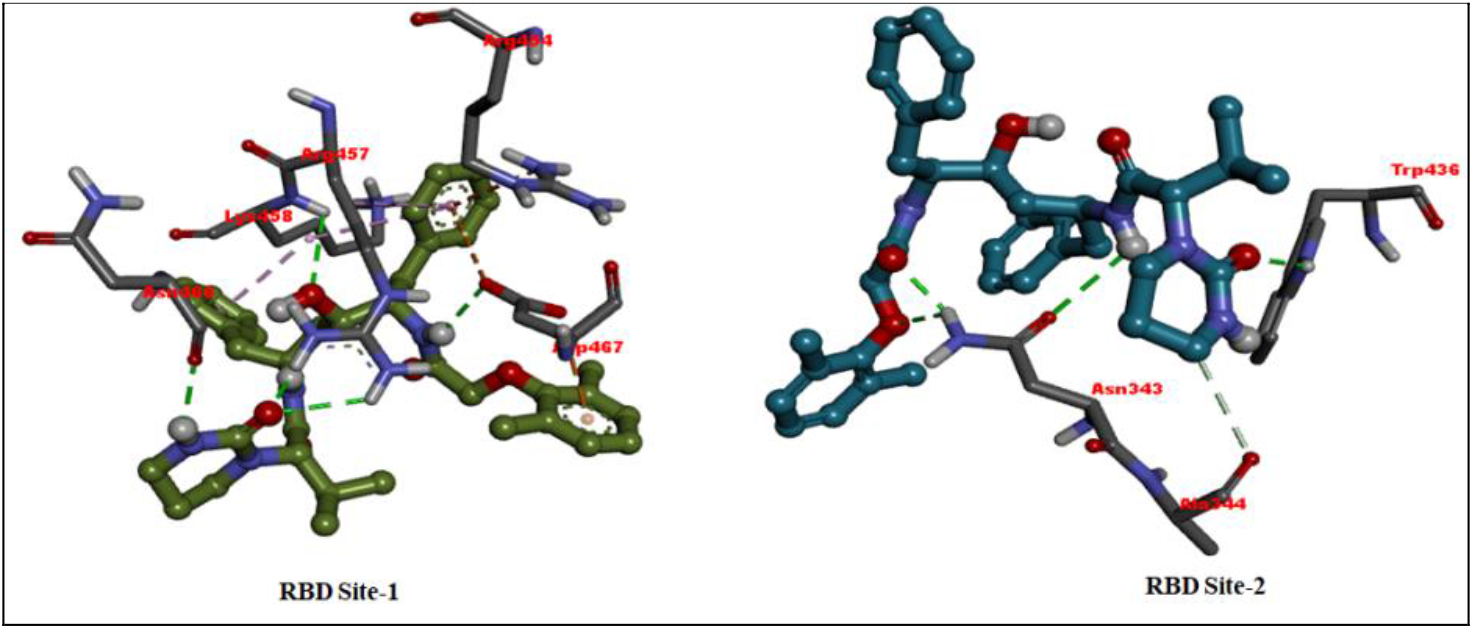
Binding orientation of Lopinavir within the active site of the RBD site-1 and site-2.

Docking analysis of Ritonavir in the site-1 revealed that it has making three strong hydrogen bonding interactions with Arg457 and another with Asp467. The drug is also involved in one hydrogen bond with Asn460 and *cation-pi* interactions with Lys458. In the binding site-2, the drug was involved in hydrogen bonding interaction with Asn343 and Trp436. The compound was found placed proximal to various hydrophobic amino acids such as Leu335, Gly339, Phe342, Ala344, Thr438, Asn437, Asn440, Ile441, Ser371, Leu368 &Phe374 and hence exhibited hydrophobic interactions. Apart from these interactions the compound was further stabilized by *pi-pi* interaction with Phe373 and Phe338.

**Figure-4:**
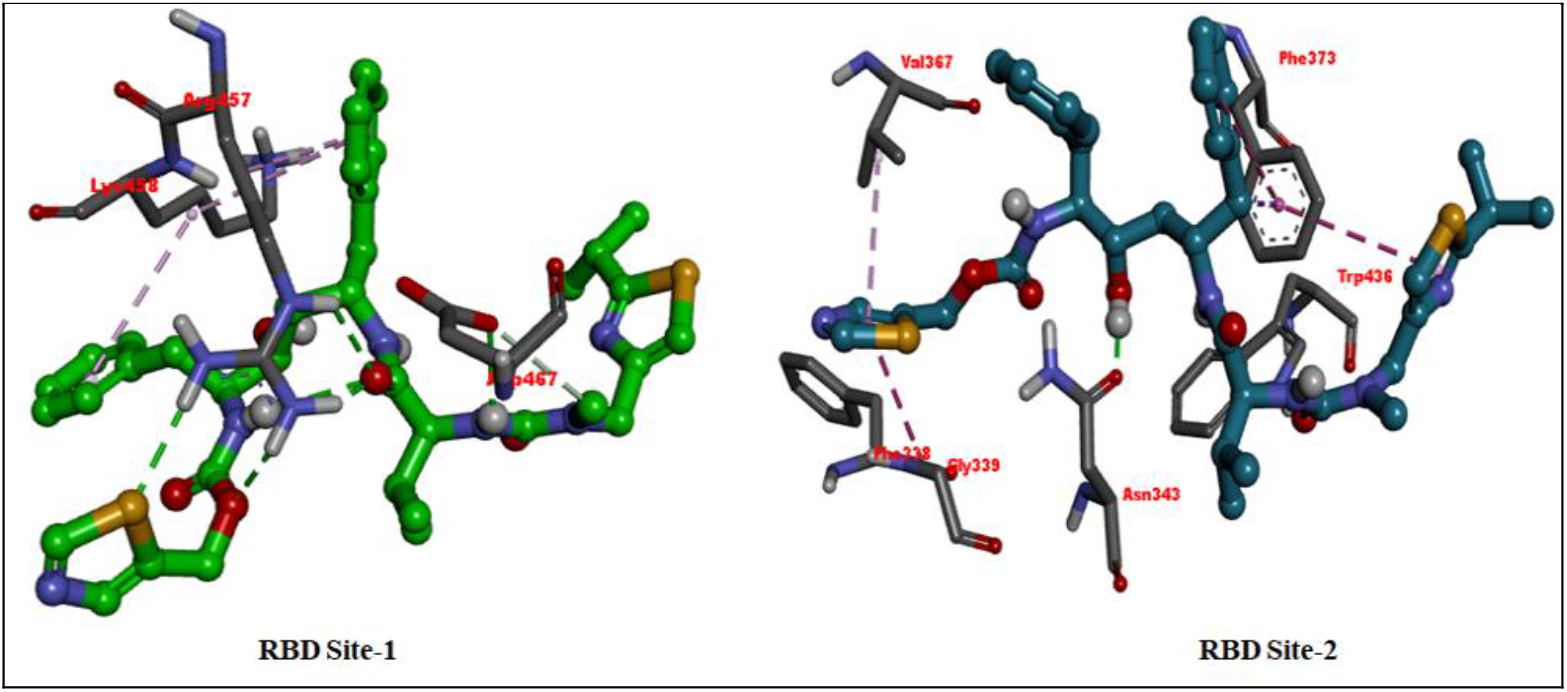
Binding orientation of Ritonavir within the active site of RBD’s site-1 and site-2.

The ligand binding analysis of the another selected drug (Astemizole) revealed a good binding affinity against the protein with the docking score of −10.1 and −10.2 in each active site of RBD domain. The drug was found to be involved in the hydrogen bon interactions with Lys458 and Gln471 in site-1 and Asn343, Arg509 in site-2 as illustrated in **Figure-5**. *In silico* analysis of this drug revealed that it has well occupied the active site.

**Figure-5:**
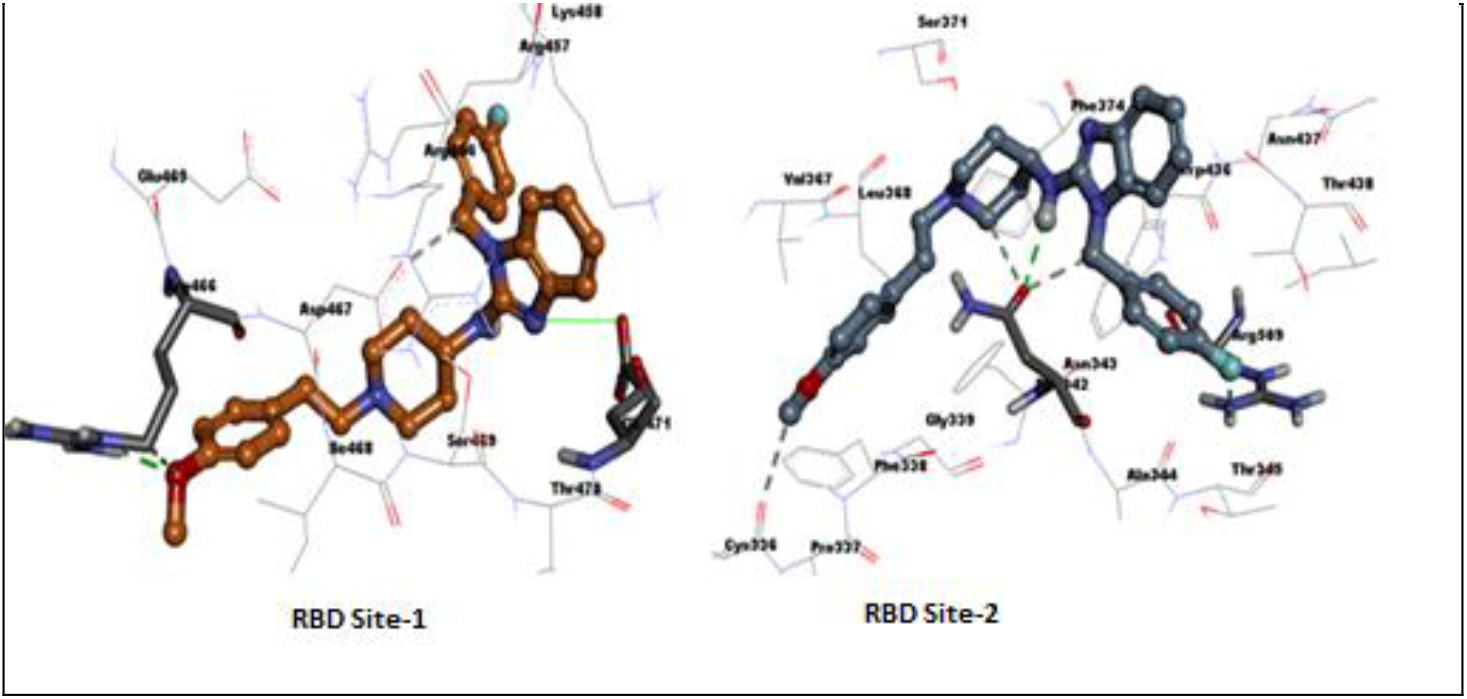
Binding orientation of Astemizole within the active site of RBD site-1 and site-2.

Amodiaquine is an aminoquinoline analog used for the therapy of malaria. Amodiaquine has been linked to severe cases of acute hepatitis which can be fatal, for which reason it is recommended for use only as treatment and not for prophylaxis against malaria. It shows good LF Rank score and ∆G in both the active sites. In site 1, the drug is making two hydrogen bonding interactions with Lys458, and Arg466. Interaction analysis in active site-2 revealed that it has hydrogen bonding interactions with Phe342, Val367 amino acid residues (**Figure-6**).

**Figure-6:**
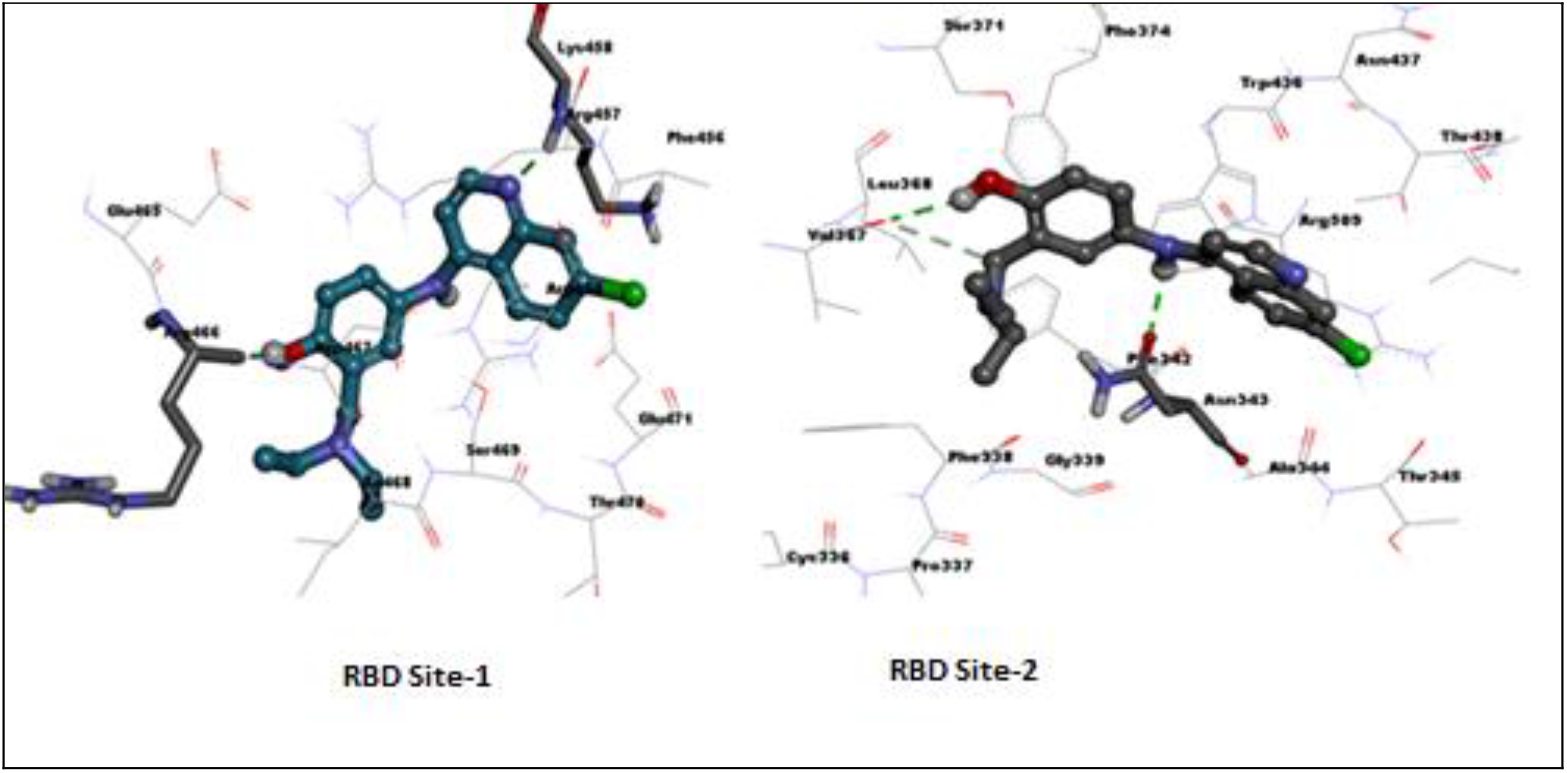
Binding orientation of Amodiaquine within the active site of RBD site-1 and site-2.

Homoharrigtonine a drug that is indicated for treatment of chronic myeloid leukemia, is the most promising drug found through *our in silico* studies. Detailed binding analysis of Homoharrigtonine revealed that it has hydrophobic interactions in the both the active sites. Docking of the Homoharrigtonine within the active site of the RBD site-1 and site-2 is illustrated in **Figure-7**. In the site-1, it showed four hydrogen bonding with Arg457, Arg466, Asp467 and Ile468 and well fitted in the hydrophobic pocket within the vicinity of Phe456, Glu465, Ser469 and Glu471. Additionally, the *cation-pi* interaction between Lys458 constituted for a stable binding profile of the drug. In site-2, the Homoharrigtonine is making hydrogen bonding with Ser371 and various non-bonded interactions. The drug is also involved in various hydrophobic interactions with Asn437, Thr438, Asn437, Thr438, Asn440, Asn343, Ala344, Thr345, Val367 and Leu368 amino acid residues.

**Figure-7:**
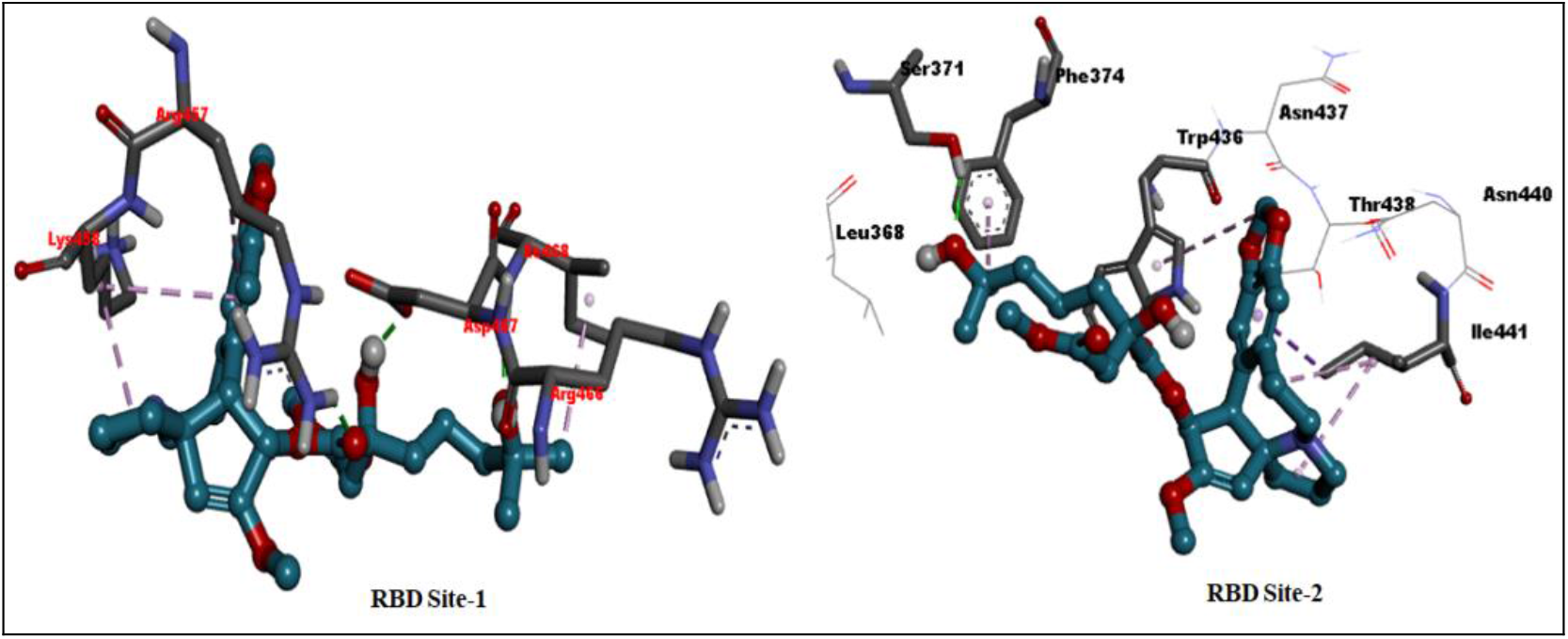
Binding orientation of Homoharrigtonine within RBD site-1 and site-2.

### 3.3 Evaluation of selected drugs in RBD-ACE-2 binding assay

In this experiment, we explored the ability of the short listed drugs on blocking binding of RBD to human ACE-2 using a commercially available competition ELISA assay. As shown below in **Figure-8,** several of the drugs showed inhibitory activity. But the stand out drug was Homoharringtonine with near complete inhibition at higher concentrations. Ritonavir and chloroquine also showed activity but the maximum activity obtained (<50%) was much less than that of Homoharringtonine. These data support further evaluation of Homoharringtonine for anti-inflammatory and anti-thrombogenic properties.

**Figure-8:**
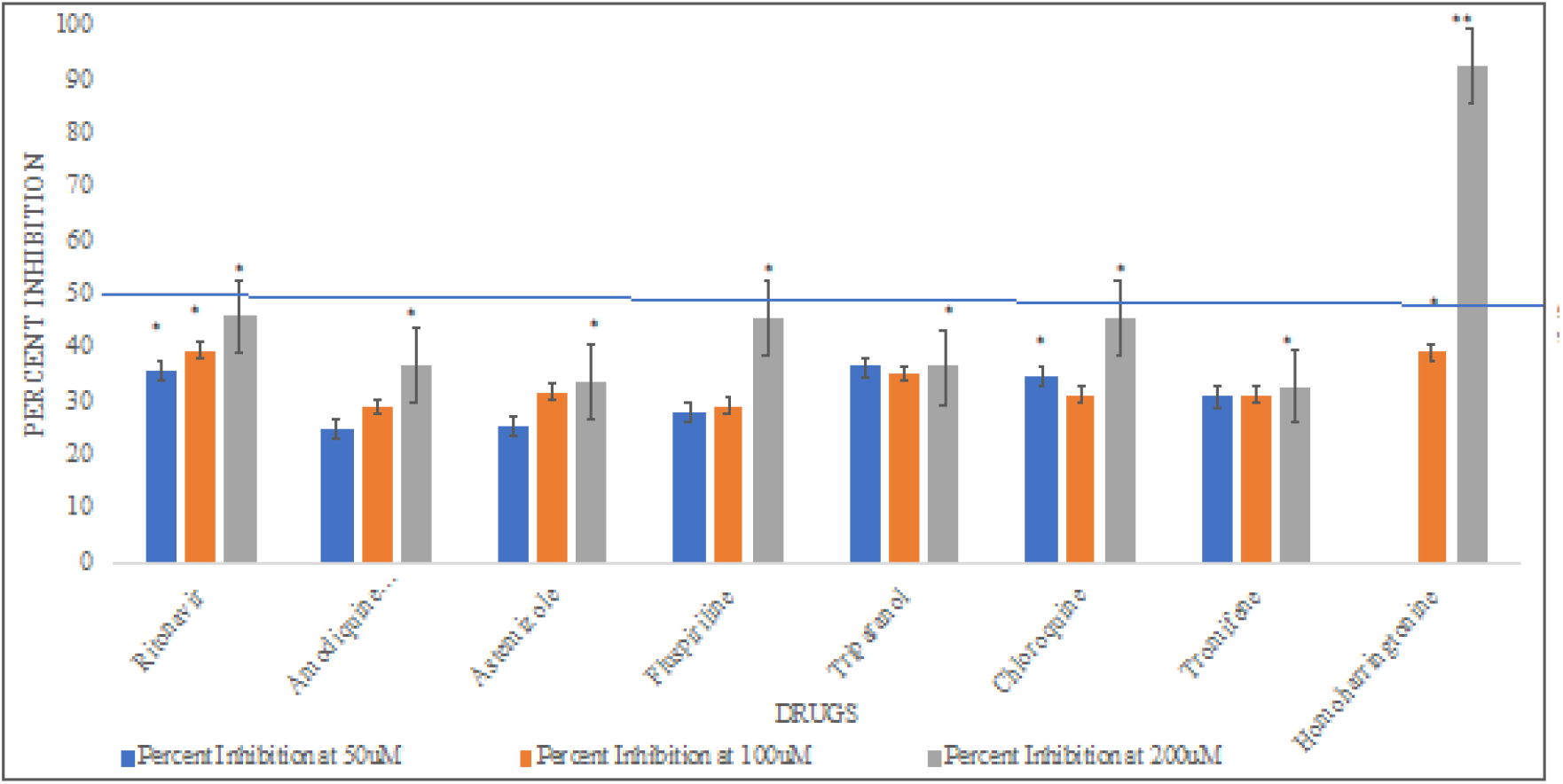
Percent Inhibition by different drugs at different concentrations of 50**μ**M, 100**μ**M and 200**μ**M. The blue line shows 50% inhibition of binding by drugs. *** P<0.001; ** P<0.01; *P<0.05. Student’s t-tests were performed to compare between the percent inhibition shown by control and drugs.

### 3.4 Testing the inhibitory effect of drugs on secretion of cytokine IL-1β

In this experiment, we explored the inherent potential of the drugs on blocking secretion of IL-1β, a cytokine known to be involved in COVID-19. We used LPS (lipopolysaccharide) as a stimulus to induce the secretion since this is a well-established inflammatory stimulant and its use does not need BLA3 safety lab unlike what is needed for using live virus. As expected LPS induced secretion of IL-1β. But none of the drugs tested showed any inhibitory activity, as shown in **Figure-9**.

**Figure-9:**
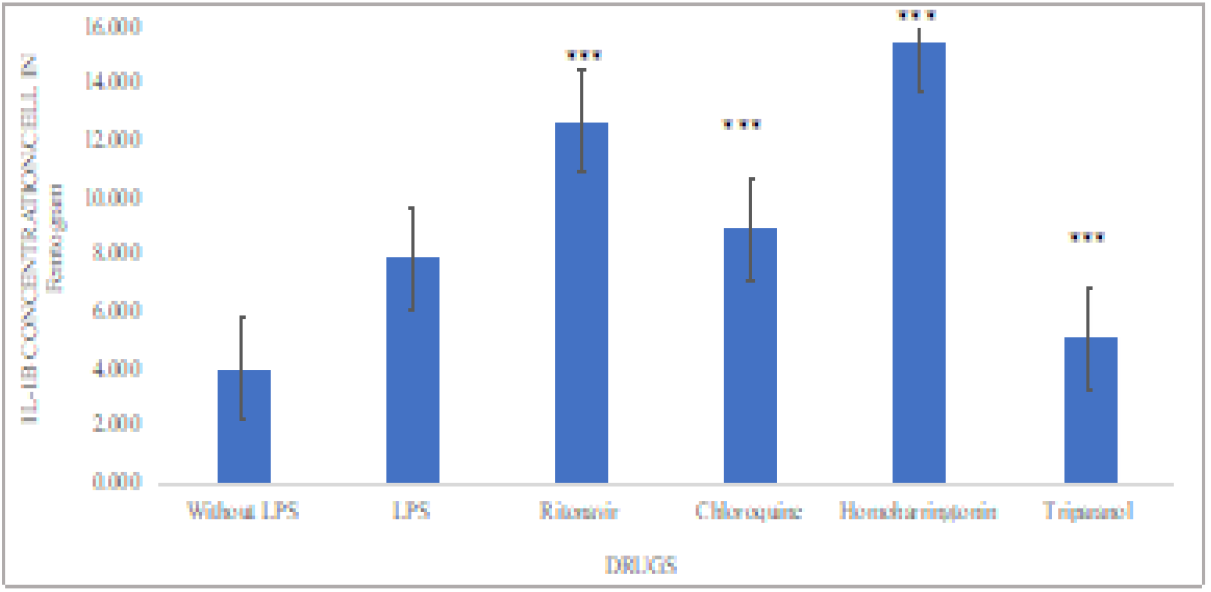
IL-1β secretion by different drugs (all drugs at 100**μ**M). *** P<0.001; ** P<0.01; *P<0.05. Student’s t-tests were performed to compare IL-1β secretion between control and LPS treated cells; and between LPS treated cells (acting as a control) and different drugs treated cells.

### 3.5 Testing the inhibitory effect of drugs secretion of Thrombomodulin

In this experiment, we explored the ability of drugs on blocking secretion of thrombomodulin, an endothelial surface protein which binds thrombin and modulates blood clotting. Inflammatory stimulus such as LPS and cytokines activate the secretion of this protein thus rendering is less functional. Secreted thrombomodulin is less effective at clotting function so drugs which reduce its secretion (therefore retaining it on endothelial surface) would be expected to have a beneficial effect. As shown below in **Figure-10**, there was induction of secreted thrombomodulin by LPS by several fold. Cytokines and LPS are known to induce secretion of surface thrombomodulin likely by activation of a protease. Of great interest was the finding that all the drugs tested showed inhibition of secretion with the best two drugs being Ritonavir and Homoharringtonine. These data are suggestive of a potentially anti-thrombogenic properties of these two drugs especially important in the context of COVID-19.

**Figure-10:**
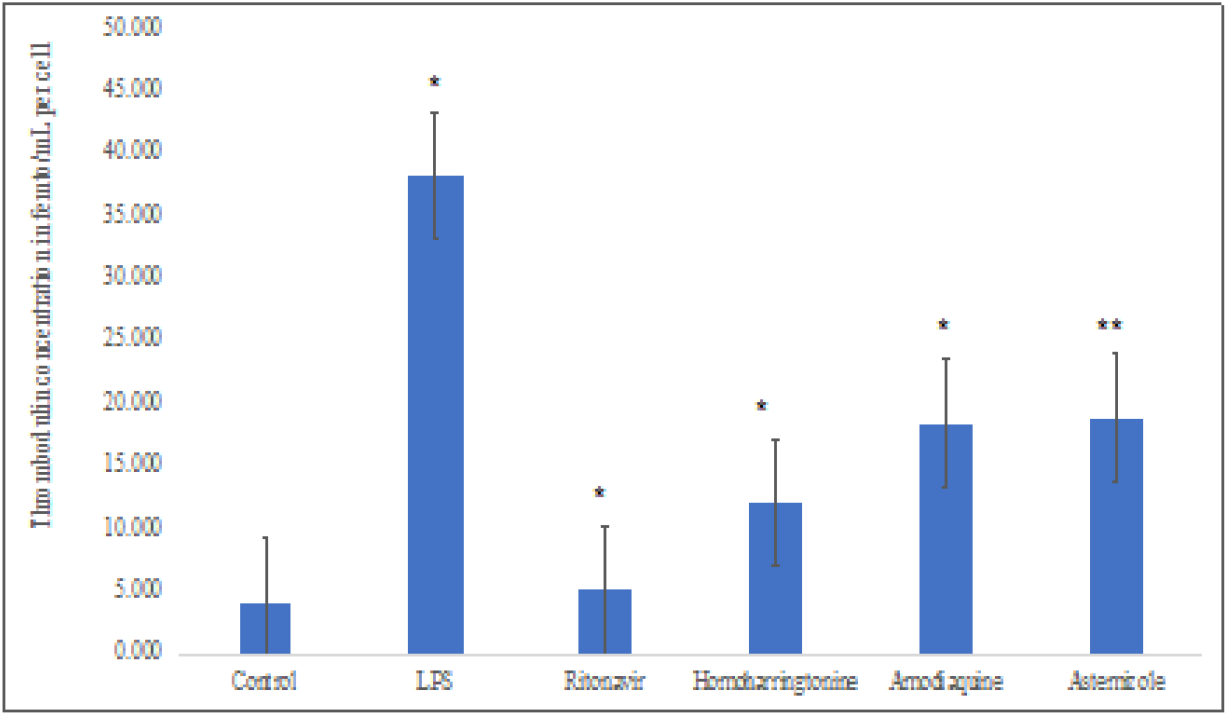
Effect of drugs Thrombodulin secretion (all drugs at 100**μ**M). *** P<0.001; ** P<0.01; * P<0.05. Student’s t-tests were performed to compare thrombodulin secretion between control and LPS treated cells; and LPS treated cells (acting as a control) and different drugs treated cells.

### 3.6 Testing the inhibitory effect of drugs on monocyte adhesion

In this experiment, we explored the ability of the drugs to reduce monocyte adhesion to the endothelium induced by inflammatory stimulus such as LPS. The adhesion of monocytes to the endothelium is the first step in the entry of these inflammatory cells into the lung during COVID-19. Thus, if a drug is able to reduce monocyte adhesion, it would be expected to be therapeutic. **Figure-11** shows there was an increase in monocyte adhesion by LPS by about 48%. While Homoharringtonine did not completely inhibit LPS induced monocyte adhesion the increase was less than that found with LPS alone (30% versus 48%). These data suggest that this drug may modulate monocyte adhesion beneficially.

**Figure-11:**
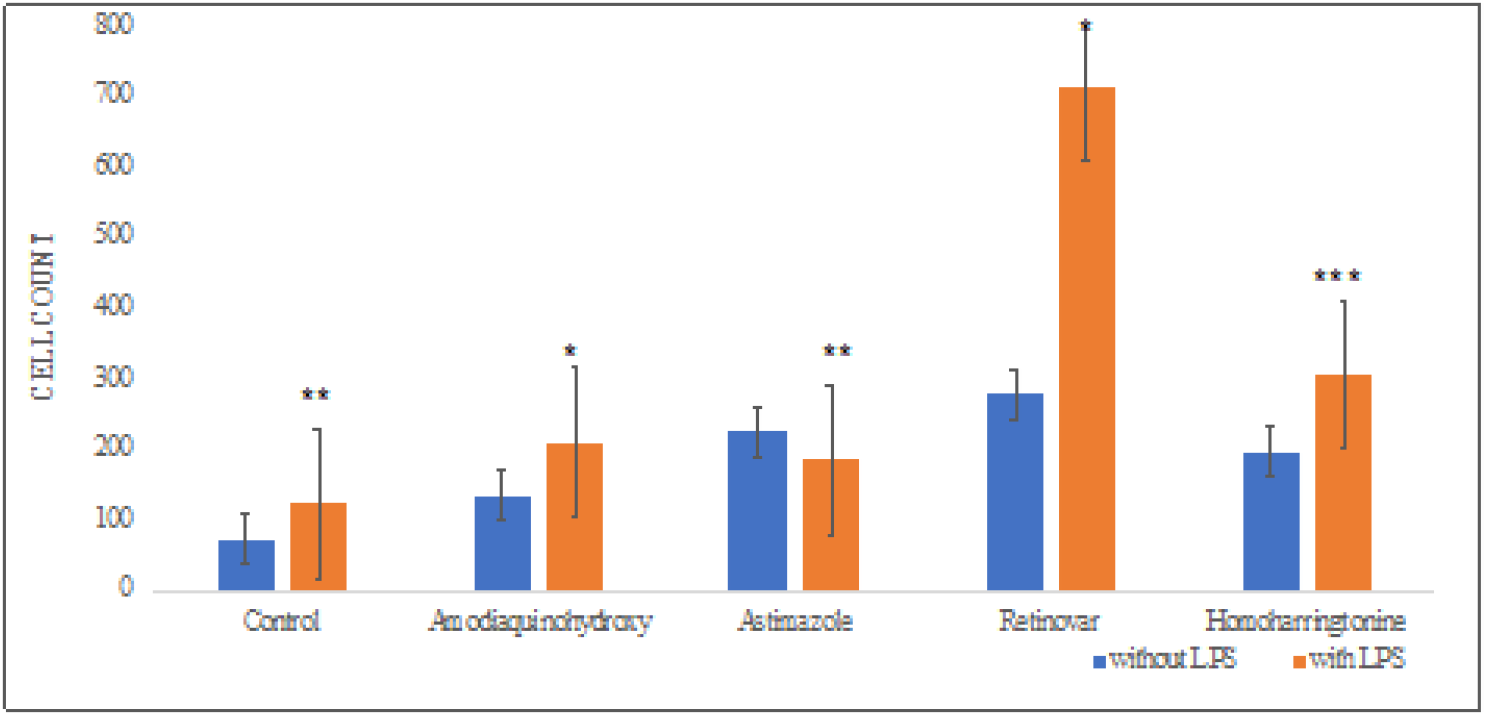
Effect on Monocyte adhesion by different drugs (at 100**μ**M) in 3D Vascular Lung Model. *** P<0.001; ** P<0.01; * P<0.05. Student’s t-tests were performed to compare between the control and drugs after treatment with LPS

## 4. Conclusion

Our work described here has identified Homoharringtonine as a potential drug candidate for repurposing for COVID-19 treatment. We find reasonable correlation between the predicted *in silico* activity of the drugs towards inhibition of RBD binding to ACE-2 with *in vitro*, suggesting that a rational drug design approach is possible for this disease. While Homoharringtonine appeared to have the greatest number of desirable properties for therapeutic intervention, Ritonavir and Chloroquine which have been tested in clinical studies showed much less activities [9]. Neither drug was able to completely inhibit binding of RBD to ACE-2 and did not possess some other properties as well. Our data would have predicted that what has been observed clinically, *i.e.* that these drugs are unlikely to be very effective in treatment of this disease (9).

Our data substantiates the observation of others with Homoharringtonine. Another report has recently shown that this drug inhibits *in vitro* replication of the virus in-Vero cells [10]. In addition, most recently it has been found to be active in a live virus induced mice model of COVID-19 [11]. The *in vivo* study reported a severe reduction in viral load upon dosing with homoharringtonine. The mechanism(s) of this efficacy were not probed in that study but in part can be attributed to its effect on viral replication and as shown by our work here potential prevention of viral entry into host cells. Since most animal models of COVID-19 represent early stage disease, our ability to demonstrate anti-thrombogenic and anti-adhesive properties in a human 3D model become very relevant in terms of the massive inflammation and micro thrombi observed in human lungs. We propose that our data with homoharringtonine using the 3D model adds to the potential mechanism by which this drug may be useful in treatment of COVID-19.

In conclusion, we propose based on our own and others work, Homoharringtonine can be tested in human trials as a nasal spray to maximize its distribution into the nasal and pulmonary tissue which are the major points of viral attachment and entry. We would also like to recognize the utility of our 3D human vascular lung model to study critical features of human disease such as thrombogenicity/clotting which is part of human disease and is also now seen as a serious adverse event with mRNA based COVID-19 vaccines.

## References

1. Dong, Yetian, et al. “A systematic review of SARS-CoV-2 vaccine candidates.” Signal transduction and targeted therapy 5.1 (2020): 1–14.

2. Kwofie, Samuel K., et al. “Cheminformatics-Based Identification of Potential Novel Anti-SARS-CoV-2 Natural Compounds of African Origin.” Molecules 26.2 (2021): 406.

3. Ehaideb, Salleh N., et al. “Evidence of a wide gap between COVID-19 in humans and animal models: a systematic review.” Critical Care 24.1 (2020): 1–23.

4. Flare, version, Cresset®, Litlington, Cambridgeshire, UK; http://www.cresset-group.com/flare/

5. Wishart, David S., et al. “DrugBank: a comprehensive resource for *in silico* drug discovery and exploration.” Nucleic acids research 34.suppl_1 (2006): D668–D672.

6. Lan, Jun, et al. “Structure of the SARS-CoV-2 spike receptor-binding domain bound to the ACE-2 receptor.” Nature 581.7807 (2020): 215–220.

7. Lead Finder, version, BioMolTech®, Toronto, Ontario, Canada; http://www.cresset-group.com/lead-finder/

8. Yavuz, Serap, and Serhat Unal. “Antiviral treatment of COVID-19.” Turkish journal of medical sciences 50.SI-1 (2020): 611–619.

9. WHO Solidarity Trial Consortium. “Repurposed antiviral drugs for COVID-19-interim WHO SOLIDARITY trial results.” New England journal of medicine 384.6 (2021): 497–511.

10. Choy, Ka-Tim, et al. “Remdesivir, lopinavir, emetine, and homoharringtonine inhibit SARS-CoV-2 replication *in vitro*.” Antiviral research 178 (2020): 104786.

11. Wen, Hai-Jun, et al. “Homoharringtonine (HHT)-A highly effective drug against coronaviruses and the potential for large-scale clinical applications.” bioRxiv (2021).

